# Vincristine treatment reverses podocyte damage in focal segmental glomerulosclerosis

**DOI:** 10.1101/2024.11.13.623397

**Authors:** William J Mason, Jennifer C Chandler, Gideon Pomeranz, Karen L Price, Marilina Antonelou, Scott R Henderson, Laura Perin, Stefano Da Sacco, Alan D Salama, David A Long, Ruth J Pepper

**Affiliations:** Developmental Biology and Cancer Research and Teaching Department, University College London Great Ormond Street Institute of Child Health, London, WC1N 1EH, United Kingdom; UCL Centre for Kidney and Bladder Health, London, United Kingdom; Royal Free Hospital, University College London Medical School, London, United Kingdom; GOFARR Laboratory, Children’s Hospital Los Angeles, Division of Urology, Saban Research Institute, Los Angeles, CA, 90027, USA; Keck School of Medicine, University of Southern California, Los Angeles, CA, 90033, USA

**Keywords:** F-actin, focal segmental glomerulosclerosis, glomerular disease, microtubules, podocytes, vincristine

## Abstract

**Introduction:** Focal segmental glomerulosclerosis (FSGS) is a significant cause of chronic kidney disease and triggered by podocyte damage which can result in cytoskeletal alterations leading to foot process effacement. Vincristine is a chemoprotective drug which alters cytoskeletal microtubules and has been used clinically to reverse FSGS. However, the mechanisms underlying the beneficial effect of vincristine are not understood.

**Methods:** We exposed immortalised human podocytes to serum obtained from an FSGS patient before, during, and after vincristine treatment. Using RNA-sequencing we determined the effect on the podocyte transcriptome alongside impacts on cytoskeletal structure and filtration barrier integrity using a glomerulus-on-a-chip model.

**Results:** We describe an adult index FSGS patient successfully treated on multiple occasions by vincristine. Podocytes exposed to serum obtained during or after vincristine treatment contained lower levels of genes associated with microtubule function compared with cells stimulated with serum collected before treatment during disease presentation. Presentation serum altered the patterning of two key podocyte cytoskeletal components, tubulin and F-actin and increased albumin permeability, changes prevented by vincristine treatment. Immunoglobulin depletion experiments revealed that the podocyte damage initiated by the presentation serum was not due to circulating autoantibodies. Defects in tubulin patterning were observed when podocytes were exposed to serum from other FSGS patients, suggestive of a common disease mechanism.

**Conclusion:** Vincristine therapy produces a milieu that protects against pathological changes induced by FSGS serum, associated with preservation of tubulin and F-actin organisation. The functional role of vincristine warrants further investigation, to advance our understanding of this alternative FSGS therapeutic.

## Introduction

Chronic kidney disease (CKD) affects approximately 10% of the global population^1^. Focal segmental glomerulosclerosis (FSGS)^2^ is associated with CKD, leading to proteinuria and ultimately progression to end stage kidney disease (ESKD). Treatment strategies such as steroids and immunosuppression are ineffective in some FSGS patients^3^. Therefore, the identification of disease mechanisms which underlie FSGS and developing new therapies to target them is needed.

The primary cause of FSGS is damage to the glomerular podocyte epithelial cells^2,3^. This can be triggered by podocyte gene mutations^4^, circulating factors^5^ or autoantibodies to slit diaphragm molecules such as nephrin^6,7^; however, in some cases the cause remains elusive. Podocytes have a unique highly branched architecture which is critical for maintaining filtration barrier integrity. This specialised shape relies on the cytoskeleton comprised of microtubules and intermediate filaments, located in the cell body and primary processes as well as F-actin, primarily located in the foot processes^8^. The F-actin and microtubule networks are intricately interlinked, through synergistic regulation of dynamics^9^, regulation of cell shape, migration and stiffness^9,10^, mitotic spindle positioning^11^, and are controlled by identical proteins, such as profilin^12^ and formin-2^13^. Disruption to cytoskeletal architecture occurs in pre-clinical *in vivo*^14^ and *in vitro*^15,16^ FSGS models as well as in cultured podocytes exposed to FSGS patient serum^17–19^. This reorganisation of the cytoskeleton changes podocyte shape resulting in cell detachment and foot process effacement^8,20^.

Several studies have tested whether therapies which preserve cytoskeletal structure improve FSGS. A small molecule targeting the GTPase dynamin, which interacts with actin, promoted stress fibre formation in cultured podocytes^21^ and improved disease progression in pre-clinical CKD models^22^. Systemic overexpression of the G-actin sequestering peptide, thymosin β-4 protected the podocyte cytoskeleton in an FSGS model of adriamycin nephropathy^15^, through the F-actin preservation. Vincristine, a chemotherapy drug^23^, which prevents microtubule polymerisation resulting in cell cycle arrest^24^ caused remission in some steroid-resistant FSGS patients^25,26^, without serious side effects^27^. These clinical findings show that vincristine is a promising therapeutic strategy for some FSGS cases. However, the mechanisms underlying the protective effect of vincristine are not fully understood. This study aimed to address this knowledge gap by collecting serum before, during, and after vincristine treatment from a steroid resistant FSGS patient. We assessed the impact of vincristine therapy on podocyte structure and transcriptomics alongside glomerular filtration barrier integrity using human cellular models.

## Materials and methods

### Patient samples

Our index patient (**Table 1**, Patient 1) was known to paediatric services^24^ with biopsy proven FSGS and subsequently presented to the adult renal clinic with a relapse of nephrotic syndrome. Serum was collected (i) before treatment when they presented with disease, (ii) during vincristine treatment and (iii) during remission whilst off vincristine treatment. To investigate the wider effect of FSGS serum on podocytes, we collected samples from three further steroid-resistant FSGS patients with significant proteinuria (**Table 2**, Patients 2-4) and age-matched healthy controls. All samples were obtained with ethical approval (05/Q0508/6 Royal Free Hospital Research Ethics Committee (REC)).

**Table 1.** Summary of treatments and clinical parameters of our index patient.

| Age | Presentation | Biopsy results | Action taken |
| --- | --- | --- | --- |
| 15 months | Nephrotic syndrome, acute kidney injury, negative immunology. | Mesangial proliferation, segmental glomerulosclerosis, nephrocalcinosis | Prednisolone (60 mg/m<sup>2</sup>/day) with no response. Cyclophosphamide (3 mg/kg/day) with partial remission |
| 2 years | Relapse | N/A | Prednisolone and cyclophosphamide gave poor response. Addition of weekly vincristine (1.5 mg/m<sup>2</sup>) caused remission |
| 7 years | Relapse | N/A | Weekly vincristine caused remission |
| 7 years 10 months | Relapse | Varying degrees of segmental hyalinosis and sclerosis | Two-month course of weekly vincristine gave partial response |
| 7 years to 11.5 years | Maintenance therapy | N/A | Vincristine every 2 weeks, reducing to every 5-6 weeks, then every 5-6 months |
| 15.5 years | Relapse | N/A | Complete response to two-month course of weekly Vincristine |
| 36 years | Relapse | N/A | Two-month course of weekly vincristine caused normal renal function in 4 months. |
| 40 years | Relapse | N/A | Two-month course of weekly vincristine caused complete remission |
| 41 years | Remission | N/A | No further action |

**Table 2.** Summary of renal parameters of each patient at the time of serum donation.

| **Patient** | **Creatinine**<br>**μmol/l** | **Albumin**<br>**g/l** | **Urine protein**<br>**creatinine**<br>**ratio mmol/l** | **Treatment at time of**<br>**sample collection** |
| --- | --- | --- | --- | --- |
| FSGS Patient 1,<br>Sample 1: relapse<br>no treatment<br>(Presentation) | 110 | 25 | 1733 | No treatment |
| FSGS Patient 1,<br>Sample 2: during<br>vincristine<br>responding<br>(Treatment) | 85 | 33 | 102 | 2 weekly vincristine |
| FSGS Patient 1,<br>Sample 3: in<br>remission off<br>treatment<br>(Remission) | 81 | 44 | 19 | Off all treatment |
| FSGS Patient 2 | 168 | 29 | 1255 | Off all treatment |
| FSGS Patient 3 | 105 | 23 | 1309 | Prednisolone and<br>cyclophosphamide |
| FSGS Patient 4 | 102 | 34 | 676 | Prednisolone and<br>ciclosporin |

### Culture of human podocytes

A conditionally immortalised human podocyte cell line was cultured as described^28^. For experiments, cells were exposed to RPMI-1640 medium, supplemented with either 10% foetal bovine serum (FBS) or patient serum for 24 hours^15^. All experiments were independently repeated four times.

### Primary glomerular cell isolation and glomerulus-on-a-chip

Human foetal kidney obtained from the Human Developmental Biology Resource (HDBR; http://hdbr.org) at 23 post conceptional weeks (PCW) with ethical approval from the National Health Service REC (23/LO/0312) was minced and digested at 37°C. The digest was flowed through 100 μm sieves three times and glomeruli collected on a 30 μm sieve, plated and grown in a monolayer. Cells then underwent magnetic activated cell sorting to isolate podocytes (using biotin anti-podoplanin antibody, BioLegend, 337015, conjugated to anti-biotin microbeads, Miltenyi Biotec, 130-090-485) and glomerular endothelial cells (gEC, using anti-CD31 microbeads, Miltenyi Biotec, 130-091-935) before expansion in VRADD (RPMI-1640, ThermoFisher, 218750910 supplemented with 10% FBS, 1% penicillin/streptomycin, 100nM 1,25(OH)2D3, 1μM all trans retinoic acid and 100 nM dexamethasone) and gEC (Cell Biologics, H1168) media respectively. We then generated the glomerulus-on-a chip in OrganoPlate 3-lane 40-well plates (Mimetas) as described^29,30^. Briefly, on consecutive days a collagen solution (5mg/ml AMSbio Cultrex 3D Collagen-I rat tail, 1M HEPES, 7g/L NaHCO_3_) matrix was firstly generated in the middle channel, before podocyte and then finally gEC seeding (20,000 of each/microchip). The plate was left for five days and then exposed for 24 hours to glomerular gEC culture media containing either 1% FBS or presentation serum, with or without 400 nM vincristine (*n = 4* microchips/condition). Medium in the capillary channel was then replaced with serum free endothelial cell media containing 40 mg/ml FITC-albumin (Merck, A9771), and the non-cellular channel medium replaced with serum-free RPMI. One hour later the urinary space media was collected and FITC-albumin absorbance read at 485nm.

### RNA sequencing

RNA was extracted from podocytes exposed to presentation, treatment or remission serums (*n = 4* repeats/condition) using the RNEasy plus kit (Qiagen). Libraries were generated with the KAPA mRNA HyperPrep Kit (Roche, KK8580) with unique dual indexes and sequenced paired-end on the Illumina NextSeq 2000 platform with an average of 16 million reads/sample (ArrayExpress accession E-MTAB-14604). FASTq files were generated and pseudoaligned to the *Homo Sapiens* genome using Kallisto for Mac^31^. Resulting count files, for each sample, were imported into Rstudio using the tximport package and the Ensembl gene IDs linked to each gene with the tx2gene() function^32^. Genes with <10 counts across all samples were removed. The DESeq2 program^33^ was used to determine differentially expressed transcripts with a statistical significance of *P*< 0.05 using the Wald test corrected for multiple comparisons between podocytes exposed to either treatment or remission serum *versus* cells stimulated with presentation serum. Resulting gene lists were ordered based on log2fold change and separated into up and downregulated transcripts.

### Tubulin and F-actin immunolabelling

Podocytes were fixed and immunolabelled as described^15^ with either Acti-Stain 488 Phalloidin (Cytoskeleton Inc., 1:140) or anti-human α-tubulin (Abcam, ab6161, 1:200) followed by goat anti-mouse secondary antibody conjugated with 594 nm fluorophore (Invitrogen, A11005, 1:200).

### Immunoglobulin G removal & Western Blot

Presentation serum was passed through NAb^TM^ Protein G Spin columns three times (ThermoFisher). Western blotting was undertaken as described^34^; samples were probed with mouse anti-human IgG antibody (1:1000, Novus Biologicals, NBP1-72758) followed by rabbit anti-mouse secondary antibodies (1:1000, Dako, P0260). Bands were detected using chemiluminescence.

### Image analysis

F-actin was analysed^15^ with the distribution in each cell determined as containing either cortical actin stress fibres accumulating around the circumference of the cells, cytoplasmic stress fibres traversing the cell body in parallel, or unorganised F-actin, showing no identifiable pattern. The tubulin organisation was classified into either being centrally nuclear concentrated or dispersed. Fifty cells were blindly analysed per condition for each independent experiment.

### Statistical analysis

Statistical analyses were performed using GraphPad Prism 10.0 for Mac. Data are presented as the mean ±SD and were assessed for normality by Shapiro-Wilk test. Statistical differences were analysed by one way analysis of variance with Tukey’s multiple comparison tests for parametric data and Kruskal-Wallis test with Dunn’s multiple comparison test for non-normally distributed data. Statistical significance was considered when *p*≤ 0.05.

## Results

### Clinical history of our index FSGS patient

Our index patient (**Table 1**) has been previously described^25^ and presented at fifteen months of age with steroid resistant nephrotic syndrome (SRNS) and a kidney biopsy demonstrating FSGS. Initially, cyclophosphamide administration led to a partial remission, but following relapse at the age of two, vincristine (1.5 mg/m^2^/week) therapy was introduced, resulting in complete remission. At the age of seven the patient relapsed and responded well to a two-month course of weekly vincristine. However, by seven years and ten months, a further relapse occurred with a second kidney biopsy demonstrating varying degrees of segmental hyalinosis and sclerosis. On this occasion, a two-month course of vincristine treatment resulted in partial remission, therefore until the age of eleven and a half the patient’s vincristine therapy was maintained, initially fortnightly then reducing to every five to six weeks, then once every five to six months. Another relapse occurred at age fifteen and a half years, with complete remission achieved by a two-month course of weekly vincristine.

After twenty years of being in sustained remission, the patient relapsed aged 36 years presenting with significant proteinuria (protein:creatinine ratio 1733 mg/mmol), hypoalbuminemia (initially 25 g/l, decreasing to a nadir of 17g/l), and acute kidney injury with serum creatinine peaking at 346 µmol/l. Weekly vincristine treatment was initiated, which resulted in a good response four months later (serum creatinine 71 µmol/l, albumin 36 g/l, urine protein:creatinine ratio 173 mg/mmol). However, following a two week pause in vincristine therapy there was a prompt relapse, resulting in re-initiation of weekly therapy and a slow wean over the subsequent five months. At this point, the patient had a creatinine of 87 µmol/l, albumin of 45 g/l and a urine protein:creatinine ratio of 55 mg/mmol. Two months later, their proteinuria was consistently less than 50 mg/mmol. A complete remission was maintained until the age of 40 when the patient relapsed once more (urine protein:creatinine ratio of 1293 mg/mmol, creatinine 77 µmol/l, albumin 27 g/l). Following a two-month course of weekly vincristine, there was a complete response with a urine protein:creatinine ratio of 26 mg/mmol, albumin 38 g/l and creatinine 78 µmol/l). At the latest consultation, just under 2 years later the patient had normal renal parameters with a creatinine of 81 µmol/l, albumin of 44 g/l and a urine protein:creatinine <20 mg/mmol.

### Vincristine treatment produces a milieu which alters podocyte microtubule genes

To understand how vincristine treatment might improve FSGS in our index patient, we examined the transcriptional profile of human immortalised podocytes^28^ exposed for 24 hours to serums (sample details in **Table 2**, Patient 1) obtained when they (i) presented with disease; (ii) were undertaking vincristine treatment; and (iii) in remission (**Fig.1a**). For each condition, 17,633 genes were detected by our analysis. Firstly, we compared podocytes exposed to either presentation or vincristine treatment serum finding 132 differentially expressed genes that were significantly (p<0.05) altered (61 upregulated, 71 downregulated in the treatment group, **Supplementary Table 1**). In contrast, only 43 genes were differentially expressed in podocytes exposed to remission *versus* presentation serum (8 upregulated, 35 downregulated in the remission group, **Supplementary Table 2**).

**Figure 1.**
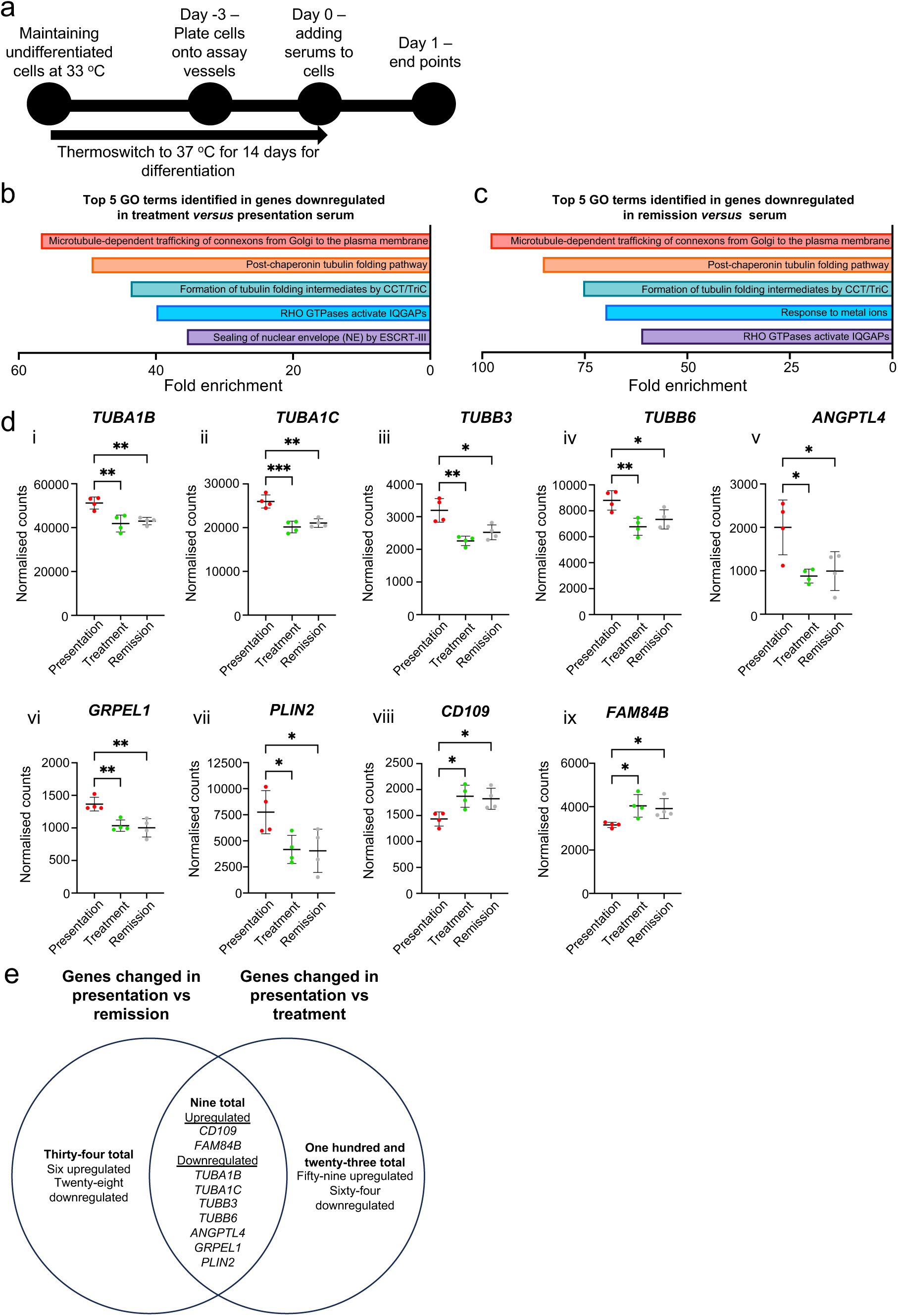
Vincristine alters the podocyte transcriptome in FSGS. **(a)** Schematic diagram demonstrating the experimental set up for all experiments. Top five gene ontology results derived from PANTHER Reactome Pathways with the input of downregulated podocyte genes compared with cells exposed to presentation serum in **(b)** cells exposed to serum taken during vincristine treatment, and **(c)** cells exposed to serum taken when the patient was in remission. **(d)** Normalised cell counts from all three treatment groups to show differential expression of **(i)** tubulin alpha 1b *(TUBA1B)*, **(ii)** tubulin alpha 1c *(TUBA1C)*, **(iii)** tubulin beta 3 class III *(TUBB3)*, **(iv)** tubulin beta 6 class VI *(TUBB6),* **(v)** angiopoietin-like 4 (*ANGPTL4),* **(vi)** GrpE like 1, mitochondrial *(GRPEL1)*, **(vii)** perilipin 2 (*PLIN2)*, **(viii)** cluster of differentiation 109 (*CD109)*, **(ix)** family with sequence similarity 84, member B *(FAM84B)*. **(e)** Venn diagram summarising differentially expressed genes analyses. Data are shown as the mean ±SD of 4 independent experiments. For PANTHER Reactome Pathway analyses, the False Discovery Rate test was used to identify significantly enriched Reactome Pathways. For individual gene plots, one way ANOVA with Tukey’s post hoc test was used. *\*p<*0.05, *\*\*p<*0.01.

We hypothesised that common biological processes were involved in both comparisons and tested this using gene ontology (GO) analyses. From the transcripts significantly upregulated when podocytes were exposed to either vincristine treatment or remission serum compared with presentation serum, no pathways were identified by GO analyses. In contrast, analysis of the downregulated podocyte transcripts, revealed that four of the five top biological pathways identified by GO analysis were identical in the cells exposed to either vincristine treatment or remission serum (**Fig.1b,c**). Three of these pathways involved microtubules or their major constituent tubulin^35^, with the fourth implicating another cytoskeletal protein^36^, Ras homolog family member A (RhoA).

We delved deeper into our gene lists and found nine common transcripts altered when podocytes were exposed to either serum obtained during vincristine treatment or remission compared with presentation serum (**Fig.1e**). In accord with our GO analysis, four of these were tubulin isoforms (*TUBA1B, TUBA1C, TUBB3* and *TUBB6*), whose levels were significantly reduced in podocytes exposed to either serum obtained during vincristine treatment or remission compared with presentation serum (**Fig.1d,i-iv**). Also, significantly downregulated were transcript levels of angiopoietin-like 4 (**Fig.1d,v**), a secreted molecule whose overexpression in rat podocytes results in proteinuria^37^ alongside GrpE like 1, mitochondrial (*GRPEL1*), a stress modulator in mammalian cells^38^ and the lipid droplet-binding protein^39^, perilipin-2 (*PLIN2*) (**Fig.1d,vi,vii**). In contrast, elevated transcript levels of genes encoding cluster of differentiation 109 (*CD109,* **Fig.1d,viii**) (CD109) and the oncoprotein^40^, family with sequence similarity 84, member 8 (*FAM84B,* **Fig.1d,ix**) were detected in podocytes exposed to either serum obtained during vincristine treatment or remission.

### Vincristine treatment alters tubulin distribution in human podocytes

As we identified that biological pathways involving microtubules and the levels of tubulin genes were altered in podocytes following exposure to serum obtained either during vincristine treatment or remission, we examined tubulin distribution using confocal microscopy. In podocytes exposed to either FBS (**Fig.2a,i**) or serum from healthy controls (**Fig.2a,ii**) for 24 hours, tubulin was concentrated in the centre of the cell. Exposure to serum taken during the time of presentation (**Fig.2a,iii**) altered podocyte tubulin distribution, leading to a dispersed cellular pattern which was not apparent when cells were stimulated with serum obtained during vincristine treatment (**Fig.2a,iv**) or remission (**Fig.2a,v**).

**Figure 2.**
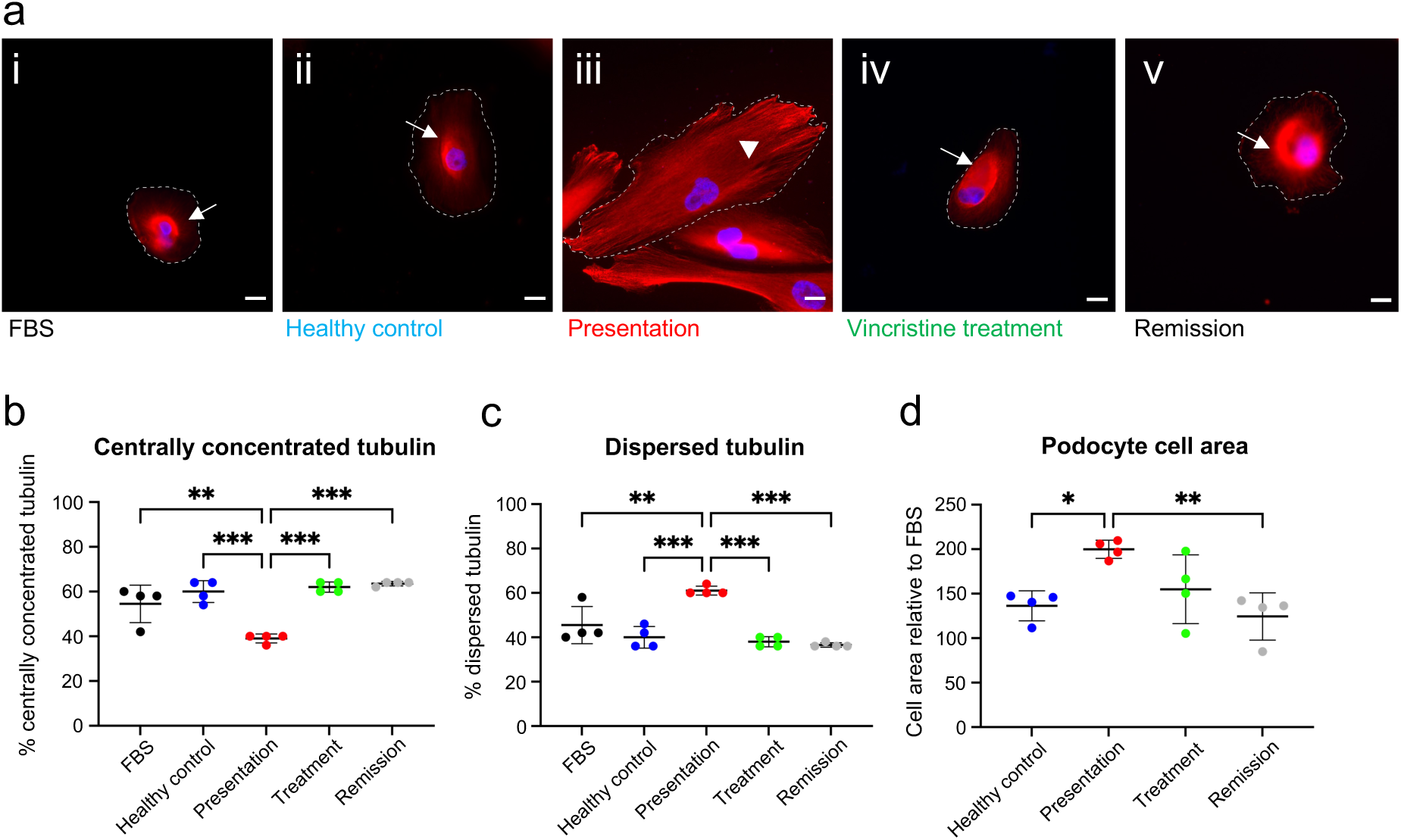
Vincristine treatment reverses changes in podocyte tubulin distribution in FSGS. **(a)** Representative images of tubulin from podocytes treated with **(i)** FBS, **(ii)** healthy control serum, **(iii)** presentation serum, **(iv)** vincristine treatment serum and **(v)** remission serum. White arrows indicate centrally nuclear concentrated tubulin and white arrowheads indicate dispersed tubulin. White dotted lines indicate cell boundaries. Scale bar = 20 μm. Red, tubulin; blue, nuclei stained with Hoechst 33342. Quantification of podocyte **(b)** centrally concentrated tubulin prevalence **(c)** dispersed tubulin and **(d)** cell area after treatment with FBS, or serum from either healthy controls or our index case during presentation, vincristine treatment or remission. Each data point represents the mean of 50 cells and data are shown as the mean ±SD of 4 independent experiments. One way ANOVA with Tukey’s post hoc test was used for all statistical testing. *\*p<*0.05, *\*\*p<*0.01, *\*\*\*p<*0.001.

We quantified the amount of centrally nuclear concentrated (**Fig.2b**) and dispersed tubulin (**Fig.2c**) in 50 cells from four independent experiments. Podocytes exposed to FBS had a centrally concentrated and dispersed tubulin prevalence of 54.5%±8.3 and 45.5%±4.3 respectively. This was similar to healthy control serum treated cells. Exposure of podocytes to presentation serum caused a shift to 39.0%±2.0 of centrally concentrated tubulin and 61.0%±2.0 dispersed tubulin (p<0.01 versus FBS, p<0.001 *versus* healthy control serum for both parameters). The prevalence of centrally concentrated tubulin was increased when podocytes were exposed to serum collected during either vincristine treatment (62.0%±2.3) or remission (63.5% ±1.0) compared with the presentation serum (p<0.001 for both comparisons). This corresponded with a reduction in podocyte dispersed tubulin (38%±2.3 and 36.5% ±1.0 for vincristine treatment and remission serum respectively) compared with the presentation serum (p<0.001 for both comparisons).

We hypothesised that the changes seen in tubulin distribution might result in alterations in podocyte size. Podocytes exposed to healthy control serum had a cell area of 136.3%±16.8 relative to FBS-stimulated cells. This rose to 199.8%±10.22 when podocytes were exposed to the serum collected during presentation, significantly higher than healthy control serum treated cells (p<0.05) and corresponded with the increase in the proportion of cells with dispersed tubulin. There was a tendency for this increase in cell size to be ameliorated when podocytes were exposed to serum obtained during vincristine treatment, with a value of 155.0%±38.6 relative to FBS, but this was not significantly different to cells stimulated with presentation serum. Finally, podocytes exposed to serum obtained during remission had a cell area of 124.4%±26.6 relative to FBS, which was significantly lower than presentation (p<0.01) serum stimulated cells (**Fig.2d**).

### Vincristine treatment results in podocyte F-actin reorganisation

Microtubules and F-actin interact in response to changes in intracellular conditions^41^. Therefore, we analysed F-actin distribution in 50 cells from four independent experiments after podocytes were exposed to: FBS (**Fig.3a,i**); healthy control (**Fig.3a,ii**); presentation (**Fig.3a,iii**); vincristine treatment (**Fig.3a,iv**); and remission serums (**Fig.3a,v**).

**Figure 3.**
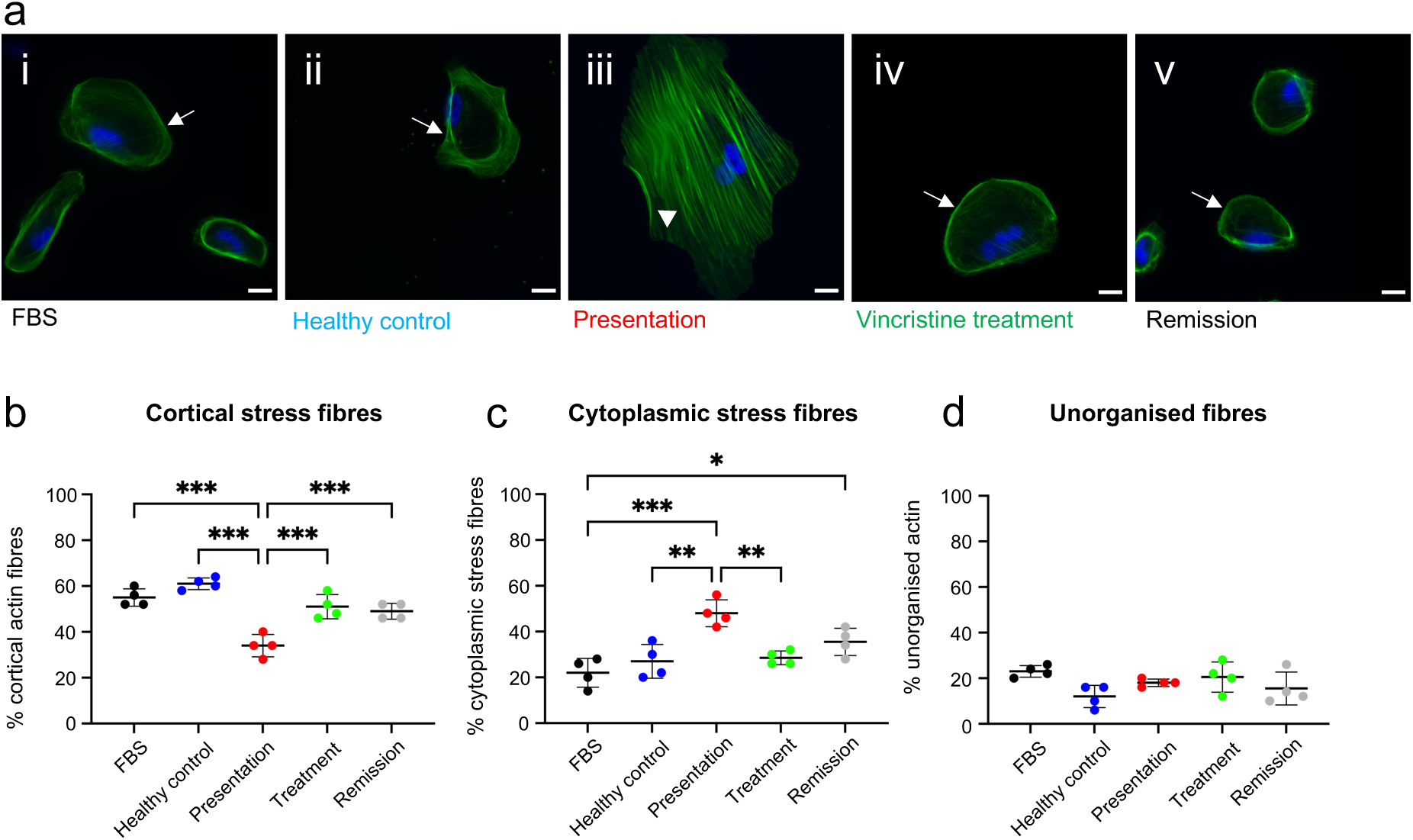
Vincristine treatment prevents podocyte F-actin reorganisation in FSGS. **(a)** Representative images of tubulin from podocytes treated with **(i)** FBS, **(ii)** healthy control serum, **(iii)** presentation serum, **(iv)** vincristine treatment serum and **(v)** remission serum. White arrows indicate cortical actin stress fibres and white arrowheads indicate cytoplasmic stress fibres. Scale bar = 20 μm. Green, F-actin; blue, nuclei stained with Hoechst 33342. Quantification of **(b)** cortical stress fibre prevalence, **(c)** cytoplasmic stress fibre prevalence, and **(d)** unorganised fibre prevalence after treatment with FBS, or serum from either healthy controls or our index case during presentation, vincristine treatment or remission. Each data point represents the mean of 50 cells and data are shown as the mean ±SD of 4 independent experiments. One way ANOVA with Tukey’s post hoc test was used for all statistical testing. *\*p<*0.05, *\*\*p<*0.01, *\*\*\*p<*0.001.

F-actin distribution was classified into cortical actin stress fibres, cytoplasmic stress fibres, or unorganised. Cells exposed to FBS, or the healthy control serum displayed 55.0%±3.8 and 61.0%±2.6 cortical F-actin respectively. Exposure to the presentation serum significantly reduced the prevalence of cortical F-actin to 34.0%±4.9 compared with both FBS and healthy control stimulated cells (p<0.001 in both cases). Podocytes exposed to the serum obtained either during vincristine treatment (51.0%±5.3) or remission (49.0%±3.5) caused a significant increase (p<0.001 in both cases) in the prevalence of cortical actin stress fibres compared with cells exposed to presentation serum (**Fig.3b**).

Podocytes exposed to FBS, or healthy control serum displayed a cytoplasmic stress fibre prevalence of 22.0%±6.3 and 27.0%±7.4 respectively. Cells exposed to presentation serum showed a cytoplasmic stress fibre prevalence of 48.0%±5.9, a significant increase compared with FBS (p<0.001) and healthy control serum (p<0.01). Exposure to the serum collected during vincristine treatment significantly reduced the prevalence of cytoplasmic stress fibres to 28.5%±3.0 compared with presentation serum treated podocytes (p<0.01). Podocytes stimulated with remission serum also displayed lower levels of cytoplasmic stress fibres (35.5%±6.0) than those apparent when cells were stimulated with presentation serum, but this was not significant (**Fig.3c**). There were no significant differences in the prevalence of unorganised F-actin between all groups (**Fig.3d**).

### Vincristine addition to presentation serum prevents albumin permeability *in vitro*

To determine if vincristine directly alters glomerular permeability, we utilised a glomerulus-on-a-chip^29^ model and performed FITC-albumin flow through experiments following addition of either FBS (**Fig.4a**), 400 nM vincristine (a dose determined from prior studies showing the maximum bolus of vincristine administered to patients is 2 mg^42^ and the average human having five litres of blood, **Fig.4b**), 1% presentation serum from the index FSGS patient without (**Fig.4c**) or spiked with vincristine (**Fig.4d**) for 24 hours (*n = 4* repeats/condition). Microchips exposed to FBS and vincristine showed no FITC-albumin flow-through into the urinary space channel, with Abs_485_ values of 0.049 ±0.00025 and 0.053 ±0.0064 respectively. Addition of 1% patient serum significantly increased the Abs_485_ to 0.49 ±0.15 (p<0.001 *versus* FBS and vincristine alone). Vincristine prevented this increase, with Abs_485_ of 0.17 ±0.15 (p<0.01 vs patient serum, **Fig.4e**).

**Figure 4.**
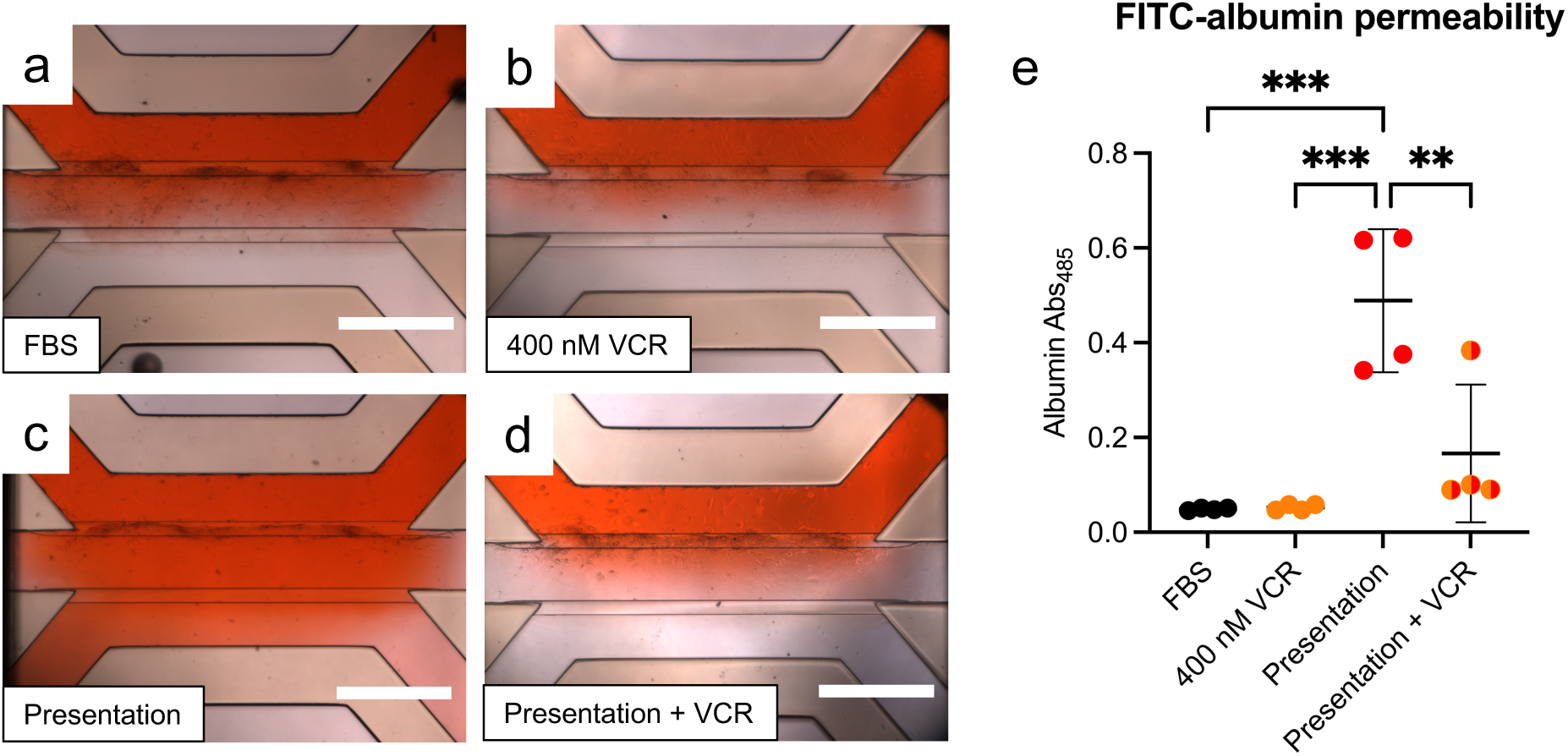
Vincristine addition to presentation serum from our index FSGS patient reduced albumin permeability. Representative images of glomerulus-on-a-chip seeded with primary human podocytes and gECs from 23 PCW kidneys exposed to **(a)** FBS, **(b)** 400 nM Vincristine, **(c)** 1% presentation serum from our index patient with FSGS and **(d)** 1% presentation serum from our index patient with FSGS spiked with 400 nM Vincristine. **(e)** Quantification of absorbance (485nm) of FITC-albumin flow through into urinary space channel. Each data point represents a glomerulus-on-a-chip with four chips used for each condition. Data are shown as ±SD. Kruskal-Wallis test with Dunn’s multiple comparisons post-hoc test were used for statistical testing. *\*\*p<*0.01, *\*\*\*p<*0.001. VCR, vincristine.

### F-actin reorganisation in our index case is not due to the presence of circulating autoantibodies

We hypothesised that the podocyte cytoskeletal reorganisation that occurred following exposure to the presentation serum in our index case could be due to circulating autoantibodies directed against slit diaphragm proteins^6, 7^. To test this, we removed IgG from the presentation serum and repeated our assessment of F-actin podocyte distribution (**Fig.5a**). Western blotting showed that three cycles using Protein G spin columns was sufficient to remove IgG from the presentation serum (**Fig.5b**).

**Figure 5.**
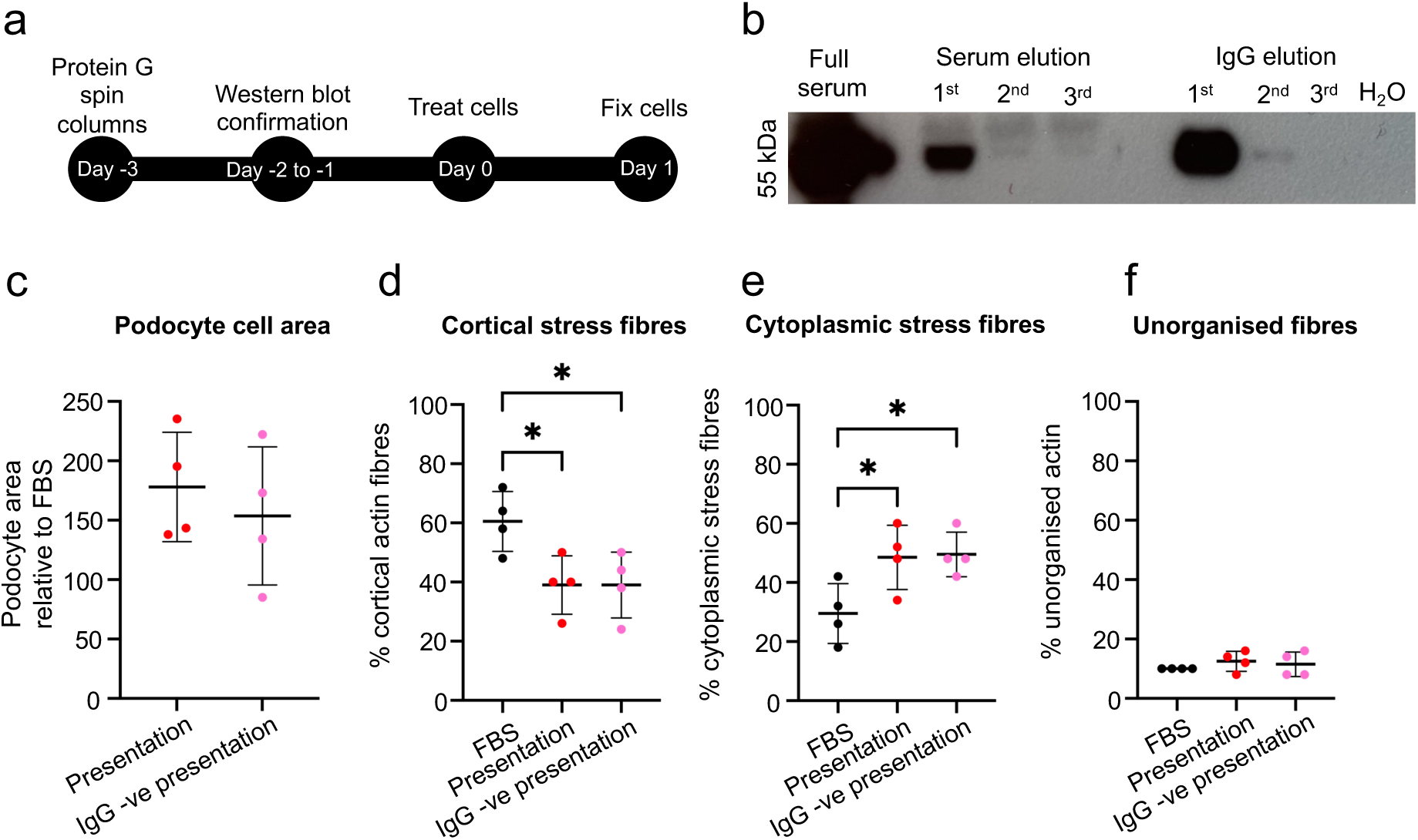
Podocyte F-actin reorganisation following presentation serum exposure was not due to IgG in patient serum. **(a)** Experimental design. Serum was depleted of IgG before treatment of human immortalised podocytes with either presentation serum or IgG depleted presentation serum. **(b)** Representative western blot of full serum, IgG depleted serum elutions and IgG elutions after 3 repetitions of Protein G spin column IgG depletion. Quantification of **(c)** podocyte cell area (unpaired *t* test). Quantification of percentage of cells displaying **(d)** cortical F-actin stress fibres (one way ANOVA with Tukey post hoc test), **(e)** cytoplasmic stress fibres (one way ANOVA with Tukey post hoc test) and **(f)** unorganised F-actin (one way ANOVA with Tukey post hoc test). Each data point shows the mean of 50 cells. Data are shown as the mean ±SD of 4 independent experiments. *\*p<*0.05.

After 24 hours of exposure, there was no significant difference in podocyte cell area (**Fig.5c**) when the cells were treated with either IgG-depleted or undepleted presentation serum (*n = 4* repeats/condition). Cells exposed to FBS showed a prevalence of 60.5%±10.1 cortical actin stress fibres, 29.5%±10.1 cytoplasmic stress fibres and 10.0%±0.1 unorganised actin fibres (**Fig.5d-f**). Podocytes exposed to undepleted presentation serum displayed reduced cortical actin stress fibres (39.0%±9.9) and increased cytoplasmic stress fibres (48.5%±10.9) compared with FBS stimulated cells (p<0.05 in both cases). Similarly, podocytes exposed to IgG-negative presentation serum had a prevalence of 39.0%±11.1 cortical actin and 49.5%±7.6 cytoplasmic stress fibres, both significantly altered compared with FBS treated cells (p<0.05 in both cases), but neither parameter was significantly different compared with the undepleted presentation serum. Unorganised F-actin was not different between any of the experimental groups.

### Serum from other FSGS patients also causes podocyte tubulin disorganisation

Finally, we tested whether exposure of podocytes to serum from three additional FSGS patients (**Table 2**, patients 2-4) also resulted in tubulin disorganisation to determine if this was a common feature of FSGS pathology. Podocytes treated with either FBS or healthy control serum displayed a similar prevalence of centrally concentrated (60.0%±7.8 and 60.0%±4.9 respectively, **Fig.6a**) and dispersed (40.0%±7.8 and 40.0%±4.9 respectively, **Fig.6b**) tubulin. The prevalence of centrally concentrated tubulin in podocytes exposed to serum taken during presentation from our index patient 1 (39.0%±2.0), patient 2 (41.5%±6.4), patient 3 (39.5%±12.3), and patient 4 (39.5%±5.5) was significantly reduced (p<0.01 for patient 1,2 and 4 and p<0.05 for patient 3) compared with cells stimulated with either FBS or healthy control serum. In contrast, the prevalence of dispersed tubulin (patient 1: 61.0%±2.0, 2: 58.5%±6.4, 3: 60.5%±12.3, 4: 60.5%±5.5) was significantly elevated (p<0.01 for patient 1,2 and 4 and p<0.05 for patient 3) compared with cells stimulated with either FBS or healthy control serum.

**Figure 6.**
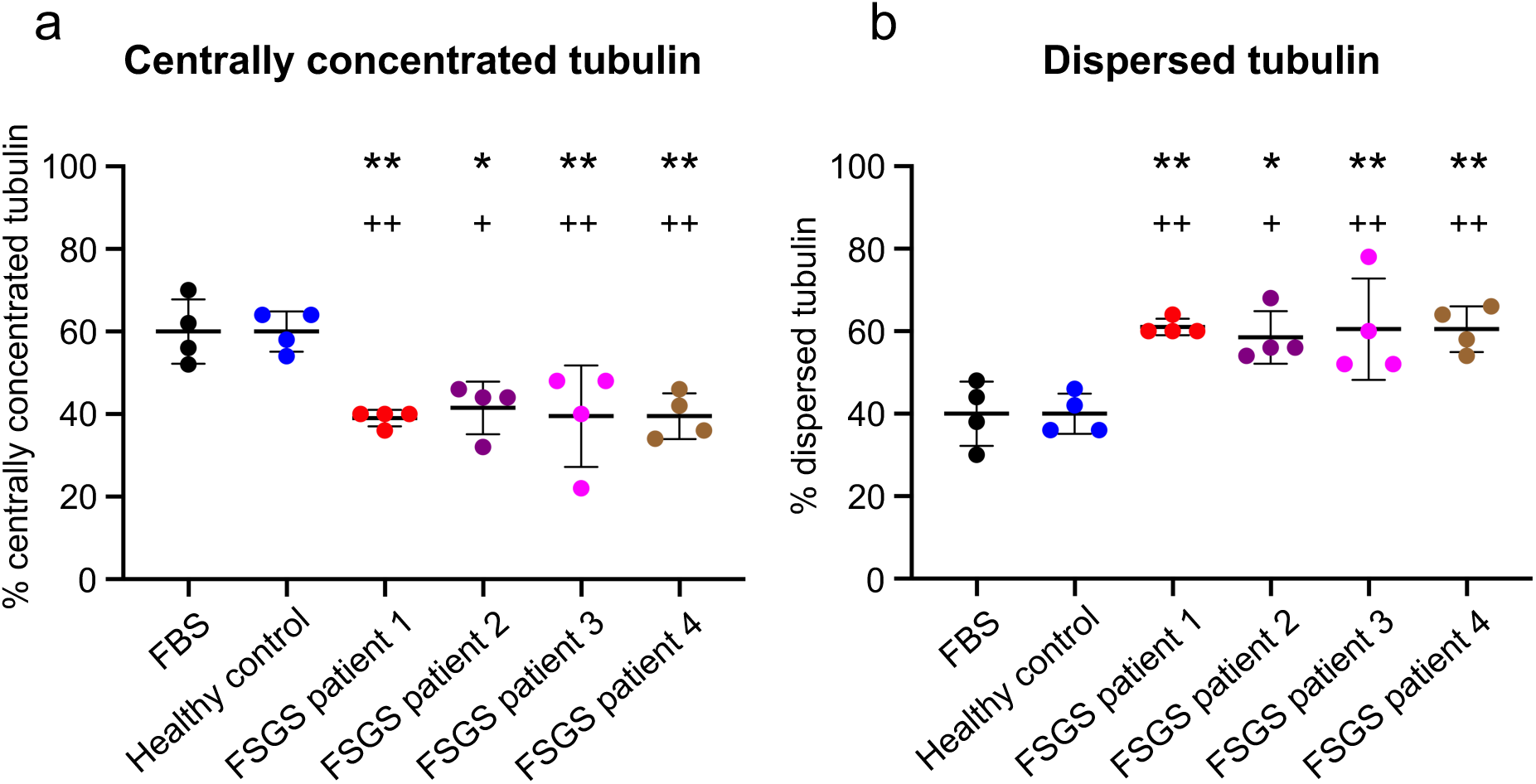
Podocyte tubulin disorganisation was found following exposure to serum from multiple FSGS patients. Podocytes were treated with 10% serum in RPMI-1640 taken from four patients (including our index case) with FSGS and the prevalence of **(a)** centrally concentrated and **(b)** dispersed tubulin quantified. FBS and serum taken from a non-renal patient were used as controls. Each data point represents the mean of 50 cells and data are shown as the mean ±SD of 4 independent experiments. One way ANOVA with Tukey’s post hoc test was used for all statistical testing. *\** - *p<*0.05 vs FBS, *\*\** - *p*<0.01 vs FBS *+* - *p<*0.05 vs Healthy control, *++* - *p*<0.01 vs Healthy control.

## Discussion

Our study outlined the cellular response to vincristine treatment for steroid-resistant FSGS in cultured human podocytes. Podocytes exposed to serum collected from our FSGS index patient during disease presentation displayed reduced levels of genes associated with microtubule function, altered tubulin and F-actin patterning alongside increased albumin permeability. These changes were prevented when podocytes were exposed to serums taken during either vincristine treatment or remission. Furthermore, we showed that podocyte tubulin disorganisation is a common feature across multiple FSGS patients. Collectively, this is the first study to link vincristine therapy with protective effects in human podocytes, in the context of FSGS.

We demonstrated that vincristine therapy can be successfully administered on multiple occasions during the relapses in FSGS to achieve a complete remission. Other studies have reported successful vincristine treatment for paediatric FSGS^25,26,43,44^. In the largest clinical trial to date, vincristine therapy for 8 weeks resulted in complete remission in 38.9%, and partial remission in 13.0% of 54 children with SRNS, without major side effects^26^. In this cohort, 32 children had biopsy proven FSGS, and 22 (68.75%) of these achieved remission after vincristine therapy^26^. To date, all publications have administered vincristine to paediatric patients. Our data suggests vincristine can also be considered for adult FSGS patients, without significant adverse effects in patients unresponsive to conventional treatments.

Reduced levels of tubulin transcripts were found when podocytes were exposed to both vincristine treatment and remission sera compared with FSGS presentation serum. Similarly, retinal pigment epithelial cells exposed to combretastatin, another microtubule destabiliser, displayed significant reductions of tubulin transcripts, a finding attributed to phosphoinositide 3-kinase (PI3K) pathway inhibition^45^. A similar mechanism may be involved in podocyte protection, a hypothesis supported by recent findings showing that mutations in the F-actin binding protein anillin that cause FSGS were associated with hyperactivation of the PI3K pathway^46^. Increased cytosol levels of β-tubulins have also been linked with reduced β-tubulin mRNA levels as an autoregulatory response in Chinese hamster ovary cells^47^. Therefore, the shift of dispersed microtubules to a centrally concentrated network by vincristine in podocytes could have increased the cytosolic concentration of tubulin, leading to a similar reduction in tubulin transcript levels.

Vincristine therapy produces a milieu which reversed the changes in microtubules and F-actin seen when podocytes were exposed to presentation serum. Cytoskeletal alterations are associated with changes to podocyte polarity, cell shape and foot process effacement^48–50^ likely resulting in the increase in FITC-albumin permeability seen with presentation serum in the glomerulus-on-a-chip model, which was prevented by vincristine. Vincristine also attenuated albuminuria and foot process effacement in a rat model of FSGS^51^. Here, the protective effect of vincristine was attributed to *α*3*β*1 integrin suppression^51^ and our RNA sequencing data also found serum collected during vincristine treatment prevented a significant increase in podocyte integrin *β*1 that was present following exposure to FSGS presentation serum (**Supplementary Table 1**). We propose that the stabilisation of both the microtubule and F-actin network by vincristine is likely to have prevented podocyte ultrastructural changes, including foot process effacement, therefore preventing albumin leakage in our index patient.

What changes to the circulating milieu might vincristine be triggering to protect podocytes? Recent studies have indicated that circulating autoantibodies against the slit diaphragm protein nephrin cause some FSGS cases^6,52–54^. Although, vincristine can have immunosuppressive effects and inhibit antibody formation^52, 53^, it is unlikely that this is the mechanism of action in our index patient, as we found no difference in the cytoskeletal changes induced by presentation serum following IgG depletion. Instead, podocyte *ANGPTL4* was significantly reduced following exposure to serum obtained during and after vincristine treatment and increased levels of this factor are associated with proteinuria in patients with diabetic nephropathy^54^, SRNS^37^ and membranous nephropathy^55^. Furthermore, transgenic overexpression of *Angptl4* in rat podocytes increased proteinuria, foot process effacement and loss of glomerular basement membrane charge^37^. Therefore, the reduction of podocyte *ANGPTL4* following exposure to treatment and remission serums from our index patient offers a possible protective mechanism of vincristine. Soluble urokinase plasminogen activator receptor (suPAR) is another proposed damaging circulating factor in FSGS^56^, which binds to podocyte plasminogen activator receptor and modulates the activity of the cytoskeletal proteins RhoA and Rac1 in a cellular model of FSGS in human podocytes^57^. Our transcriptional results show elevated *PLAU* (urokinase-type plasminogen activator) levels, the secreted natural ligand for uPAR, in cells exposed to presentation serum compared with the treatment serum; and this warrants further investigation. An intriguing alternative explanation is that vincristine directly affects podocytes, and this is suggested from our data showing that direct vincristine addition attenuates the albumin leakage seen following exposure of the glomerulus-on-a-chip to FSGS presentation serum from our index patient.

In conclusion, we provide the first evidence that vincristine therapy produces a milieu that protects against pathological changes induced by FSGS serum, associated with the preservation of tubulin and F-actin organisation. Furthermore, we demonstrate the effectiveness of vincristine in both paediatric and adult steroid resistant FSGS, warranting further investigation to advance our understanding of this alternative FSGS therapeutic.

## Disclosure

The authors have nothing to disclose

## Acknowledgements

We thank UCL Genomics for performing RNA-sequencing. We thank the HDBR, funded by Medical Research Council (MRC) and the Wellcome Trust. This work was supported by a Wellcome Trust Investigator Award (220895/Z/20/Z, to DAL), a Kidney Research UK (KRUK) Intermediate Fellowship (INT_004_20210727 to JCC) and a donation from the family of a patient with FSGS. The Long laboratory is supported by the National Institute for Health Research (NIHR) Biomedical Research Centre at Great Ormond Street Hospital for Children NHS Foundation Trust who also provided a Travelling Fellowship and Small Project Grant to WM for this work and the LifeArc-Kidney Research UK Translational Centre for Rare Kidney Disease.

## Author Contributions

WJM, JCC, DAL and RJP conceived and designed the project. WJM performed experiments. WJM, JCC, GP and DAL analysed and interpreted the data. KLP and LP and SDS provided reagents and technical expertise for the glomerulus-on-a-chip assay. DAL, ADS and RJP supervised the study. WJM collated and presented the figures and WJM, JCC, DAL and RJP wrote the paper which all authors approved.

## Notes

### Competing Interest Statement

The authors have declared no competing interest.

